# Lipid droplet structural remodeling in adipose tissue upon caloric excess

**DOI:** 10.1101/2021.08.23.457409

**Authors:** Weinan Zhou, Sarith R Bandara, Cecilia Leal, Sayeepriyadarshini Anakk

**Affiliations:** Department of Molecular and Integrative Physiology, University of Illinois at Urbana-Champaign, Urbana, Illinois 61801, United States; Department of Materials Science and Engineering, University of Illinois at Urbana-Champaign, Urbana, Illinois 61801, United States; Cancer Center at Illinois, University of Illinois at Urbana-Champaign, Urbana, Illinois 61801, United States

**Author notes:** These authors contributed equally to this work (Weinan Zhou, Sarith R Bandara). Corresponding authors. (C.L.); (S.A.).

## Abstract

Excess calories are stored as triacylglycerols (TAG) and cholesteryl esters (CE) in lipid droplets (LD), and during obesity, LD expansion occurs. X-ray scattering of adipose tissue uncovered that LDs comprise two TAG packing domains: a disordered core and a multilamellar shell. The number of TAG layers increases upon diet-induced obesity and is adipose depot-specific. Further, collagen was highly oriented in brown but randomly dispersed in white fat. We discovered that the body’s surfactant, bile acids (BAs) stimulate remodeling of LD size. Deleting the BA receptor, Farnesoid X receptor (FXR) reduced a hydrophilic BA, β muricholic acid (β-MCA), and enlarged the adipocytes. BA composition is a critical determinant of overall hydrophobicity index and solubilization ability. Accordingly, we found that the obesogenic diet reduced a hydrophobic BA, chenodeoxycholic acid (CDCA). Taken together, these findings implicate that BAs, tissue niches, and diet influence LD structural remodeling.

**Summary:** Lipid droplets (LDs) pack triacylglycerols (TAGs) with altered dimensions and exhibit distinct collagen orientation between the white and brown fat depots and are remodeled by bile acids (BAs) such that deletion of BA-receptor, Farnesoid X receptor (FXR) results in adipocyte hypertrophy.

## Introduction

Obesity is the second leading cause of preventable death in the United States (*1*). Adipose tissue routinely expands during obesity to store excess calories within lipid droplets (LDs) in the form of triacylglycerols (TAGs) and cholesteryl esters (CEs) (*2*). Rapid and prolonged LD expansion is strongly associated with metabolic diseases. However, the biophysical and genetic mechanisms controlling LD remodeling during abundant nutrition remain elusive.

LDs are typically represented as droplets containing disordered TAGs and CEs; yet, how fat structures itself inside the LDs is not fully understood. LDs must efficiently pack fat while nimbly remodeling and expanding in response to metabolic alteration. Therefore, the molecular structural organization in LDs likely contributes to this process. Recently, Shimobayashi and Ohsaki (*3*) demonstrated that LDs in human hepatocarcinoma cells appear to organize fat in a layered structure at their periphery, and that their organization alters according to the cellular state (*4*). Previously, in human osteosarcoma cells (*5*), a model of layered LDs was postulated where TAGs were surrounded by liquid crystalline (LC) domains of CEs, with the domain size dependent on the TAG:CE ratio. Thus, we anticipated that the LDs of adipocytes (cells of primary fat storage) will remodel their molecular organization to optimize packing, especially in obesity. In this report, we examined fat packing and LD structural properties under normal and obesogenic diets in the different adipose depots.

Bile acids (BAs) are natural detergents of the body known to emulsify fat and form micelles to promote solubilization and absorption (*6*). Recently, BA-loaded microparticles were proposed as an alternative to invasive liposuction for the removal of undesired fat deposits (*7*). Moreover, the role of BAs as signaling molecules that activate cell surface and nuclear receptors has been well established (*8, 9*). Nuclear receptor, Farnesoid X receptor (FXR), a major endogenous BA receptor, is expressed in many tissues (*10, 11*) and transcriptionally controls lipid metabolism (*12, 13*). Previous studies indicated an *in vitro* role for FXR in the differentiation of white adipocytes (*14, 15*), and recent work revealed that overexpression of human *Fxr* in mice increased white adipocyte size (*16*). We conjecture that BAs will impact LD size, and that the BA-FXR axis may control LD remodeling in adipocytes.

Using X-ray scattering, laser scanning confocal microscopy (LSCM) and a novel adipocyte-specific *Fxr* knockout (Ad-*Fxr*KO) mouse model, we examined how diet, BAs, and FXR affect the biophysical properties of LDs in normal and obese conditions.

## Results and Discussion

### Deletion of adipose *Fxr* expression results in adipocyte hypertrophy

FXR is expressed in white adipocytes (*11*) and is known to regulate fat metabolism (*12, 13*). We investigated and found detectable levels of *Fxr* transcripts in mature adipocytes isolated from both white adipose tissue (WAT) and brown adipose tissue (BAT) with higher mRNA levels being present in WAT (fig. S1A). To define the role of FXR in adipocytes, we generated Ad-*Fxr*KO mice and confirmed *Fxr* knockdown by qRT-PCR analysis (fig. S1B). We then challenged Ad-*Fxr*KO and f/f *Fxr* mice with either chow, 60% high-fat diet (HFD), or western diet (WD) for 4 weeks. As expected, HFD- and/or WD-fed mice showed increased body weight and fat mass compared to chow-fed mice (fig. S1, C and D), which correlated with the increase in transcripts of key lipid synthesis genes *Pparg* and *Dgat2* in WAT (fig. S2B) and BAT (fig. S3B), while SE synthesis genes *Soat1* and *Soat2* were elevated in WAT only after the diet (fig. S2C). The lipolytic gene *Hsl* was decreased in WAT (fig. S2A) but increased in BAT (fig. S3A) upon HFD, indicating an inverse regulation between the two depots. This decrease in WAT *Hsl* expression is not dependent on FXR (fig. S2A). Another lipase, *Atgl* transcript was increased in HFD- and/or WD-fed conditions compared to the normal chow distinctly in BAT (fig. S3A). Despite these gene changes, Ad-*Fxr*KO and f/f *Fxr* mice exhibited similar weight gain and fat mass except for BAT, which weighed higher in WD-fed Ad-*Fxr*KO than WD-fed f/f *Fxr* mice (fig. S1, C and D).

We then investigated if there is a difference in individual adipocyte size in the presence and absence of FXR expression. Ad-*Fxr*KO mice showed a notable adipocyte hypertrophy specifically in the white adipose under normal diet (Fig. 1, A and B and fig. S4A). When challenged with obesogenic conditions, control animals revealed the expected increase in the size of white and brown adipocytes (Fig. 1, A to D and fig. S4, A and B). Intriguingly, Ad-*Fxr*KO mice displayed adipocyte hypertrophy in both fat depots compared to the f/f *Fxr* mice (Fig. 1, A to D and fig. S4, A and B) under HFD and WD. A recent study exhibits a contrasting observation that overexpression of human FXR in mouse adipose tissue enlarges white adipocytes and limits their capacity to expand during obesity (*16*). A caveat to the ectopic *Fxr* expression study is that the 3- to 5-fold overexpression is driven by aP2 promoter (*16*), which can be leaky and induce *Fxr* expression in brain, heart, skeletal muscle, endothelial cells, and adipose-resident macrophages in addition to adipocytes (*17, 18*), while our findings are based on the analysis of adipocyte-specific *Fxr* knockout (Ad*Fxr*-KO) mice. We also found that Ad-*Fxr*KO mice exhibited a reduction in the expression of some of the lipogenic and lipolytic genes in white (fig. S2) and brown (fig. S3) fat depots compared to the f/f *Fxr* mice. These findings suggest that adipose FXR may transcriptionally regulate lipid metabolism in adipose tissues.

**Figure 1.**
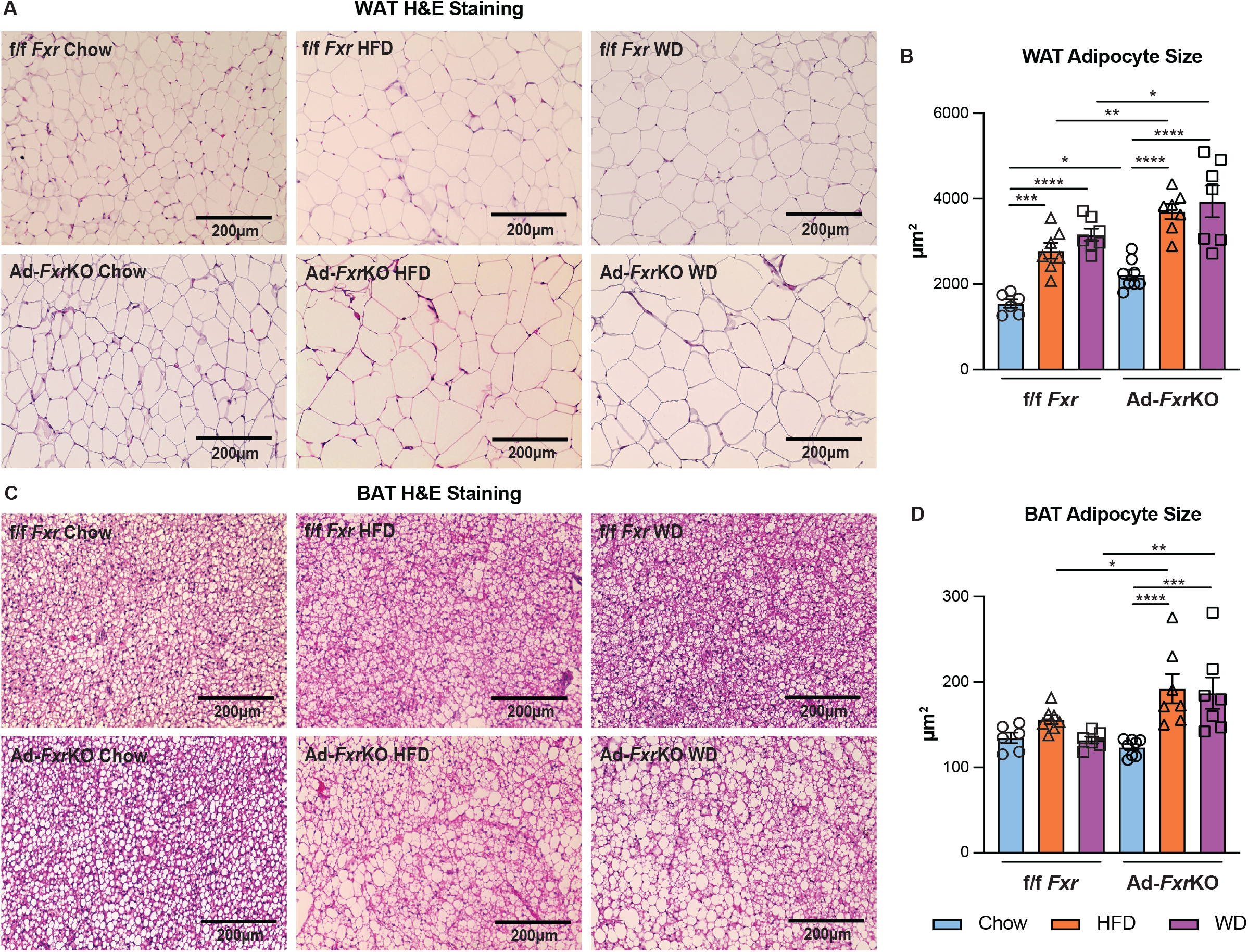
Deletion of *Fxr* results in adipocyte hypertrophy. (A-B) Representative images (A) and the mean (B) of adipocyte size of H&E-stained WAT sections of Ad-*Fxr*KO and f/f *Fxr* mice upon different diets for 4 weeks (n=6-8 mice per group). (C-D) Representative images (C) and the mean (D) of adipocyte size of H&E-stained BAT sections of Ad-*Fxr*KO and f/f *Fxr* mice upon different diets for 4 weeks (n=6-8 mice per group). Data are represented as mean ± SEM. \**P* < 0.05, \*\**P* < 0.01, \*\*\**P* < 0.001, \*\*\*\**P* < 0.0001.

### Diet affects fat packing in an *Fxr*-independent manner

LDs are often treated as amorphous droplets of neutral lipids—predominantly TAGs and CEs— stabilized in the aqueous cytosol by a lipid monolayer and proteins. However, a few recent reports have uncovered that LDs, in cancer or mitotically arrested cell lines, may adopt a liquid crystalline (LC) structure where TAGs and CEs self-organize in layers at an average separation of 3-5 nm (*3-5*). Adipocytes are challenged to store a lot of fat during obesogenic conditions and a LC structure in LDs is a much more efficient way to pack TAGs and CEs than an amorphous configuration.

Synchrotron small-angle X-ray scattering (SAXS) data of both brown and white adipose tissues from Ad-*Fxr*KO and f/f *Fxr* mice fed different diets displayed diffraction peaks at *q*^001^= 0.15, *q*^002^=0.30, and *q*^003^=0.45 Å^−1^, which are consistent with the presence of a multilamellar liquid crystalline structure with inter-lamellae separation *d*^001^=2π/*q^001^* = 42 Å (Fig. 2A). The multilamellar structure is rather robust persisting to temperatures up to 57 °C (fig. S5). This repeat spacing exceeds by 5 Å of what is expected for CE layers but closely matches the spacing of TAG layers (*19*) (inset in Fig. 2C). The different possible conformations of TAGs within each layer is depicted in fig. S6 (*20*).

**Figure 2.**
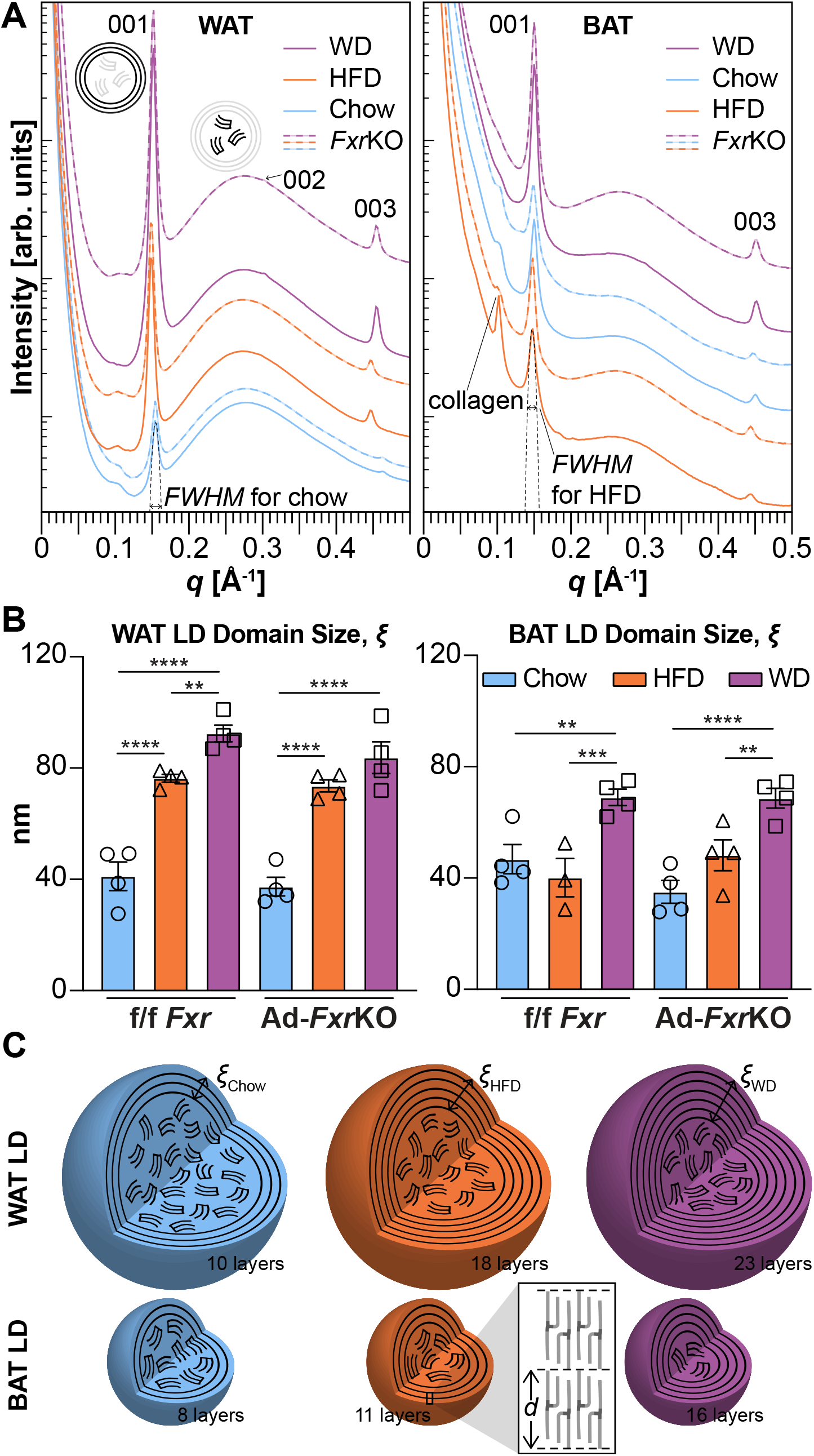
Diet affects fat packing within lipid droplets in adipose tissue in an *Fxr*-independent manner. (A) SAXS data of BAT and WAT from Ad-*Fxr*KO and f/f *Fxr* mice fed different diets display a set of three peaks in the *q range* of 0.15, 0.30, and 0.45 Å^−1^ that arise from (001), (002), (003) diffraction of multilamellar TAG layers in LDs. Multilamellar peaks coexist with a broad peak arising from a disordered domain of TAGs and CEs (n=4 mice per group, 20 locations on tissue per mouse). (B) X-ray data estimation of the size of the layered domains (in nm) obtained for different depots, diets, and Ad-*Fxr*KO *vs* f/f *Fxr* mice (n=4 mice per group). (C) The number of multilamellar layers in an LD depends on diet. WD yields LDs with the largest number of fat layers. The magnified inset shows how TAG molecules are arranged within the lamellae.

In coexistence with the characteristic multilamellar (*h,k,l*)=(001), (002), and (003) peaks, there is a broad peak at *q* = 0.27 Å^−1^ that is consistent with the presence of a disordered fat domain (*21–24*). This is in line with an LD structure where fat packs in a multilamellar LC configuration towards the LD rim and is disordered at the core (*4*). How TAGs and CEs partition into the disordered and layered domains is still an open question. CEs and TAGs could phase-separate into either domain (*4, 5*) or mix at the molecular level and distribute into the disordered and layered regions. A peak at lower *q* = 0.1 Å^−1^ is one of the diffraction peaks arising from a periodic arrangement of collagen fibers in the tissue extracellular matrix (ECM) which is generally present at *q* between 0.05 and 0.15 Å^−1^ (*25*). While the multilamellar peak position, and hence the spacing between fat layers, appears constant for both adipose tissues and diets, there is a noticeable difference in peak intensity and width. This is important because peak full-width at half maximum *FWHM* directly relates to domain size (*26*) 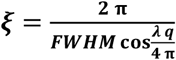 *i.e.* how many layers of fat are packed within a LD (Fig. 2B). For example, WAT LDs have a domain size of 76.4 nm and a layer-to-layer distance of 4.2 nm, which means that there are on average 18 repeating layers of TAG in WAT *vs* 23 layers under WD. This suggests that in response to dietary challenges, LDs remodel not only size, but structure or number of fat layers (Fig. 2C). Excess nutrients from WD and HFD result in lower CE/TAG ratio and an increase in the domain size, which is consistent with TAG being enriched in the layered domain.

WD results in LDs with the highest number of fat layers in both WAT and BAT. In WAT, HFD yields LDs with more layers than chow, but no significant difference is observed in BAT. The main difference between HFD and WD is the presence of sucrose in WD. We conjecture that sucrose may facilitate the packing of TAGs in bigger LDs with more packed layers via an osmotic effect. No significant difference in domain size was observed between Ad-*Fxr*KO and f/f *Fxr* genotypes (Fig. 2B).

### Adipose depots display distinct collagen orientation

Remodeling of the ECM is associated with and contributes to adipose expansion (*27*). Collagen is the largest group of ECM proteins (*28*) and often yields a diffraction signal because fibers are bundled at specific and periodic distances from each other (*29*). Two-dimensional diffraction patterns of collagen in adipose tissue (Fig. 3A) revealed a stark asymmetric signal for BAT, indicating that the ECM collagen fibers are highly oriented when compared to a symmetric diffraction ring indicating random collagen distribution in WAT. This result is consistent with histology data with collagen stained with picrosirius red (Fig. 3, B and C). Collagen has a pericellular deposition in WAT (Fig. 3B) and reticular deposition in BAT (Fig. 3C). Additionally, a more intense collagen peak at *q* = 0.1 Å^−1^ was observed in the brown compared to white adipose depot (Fig. 2A), supporting that collagen is well oriented and better packed in BAT compared to WAT. Additional experiments are necessary to fully understand the role of collagen morphology in obesity, but we infer that collagen orientation may restrict LD growth and remodeling in an adipose-depot specific manner.

**Figure 3.**
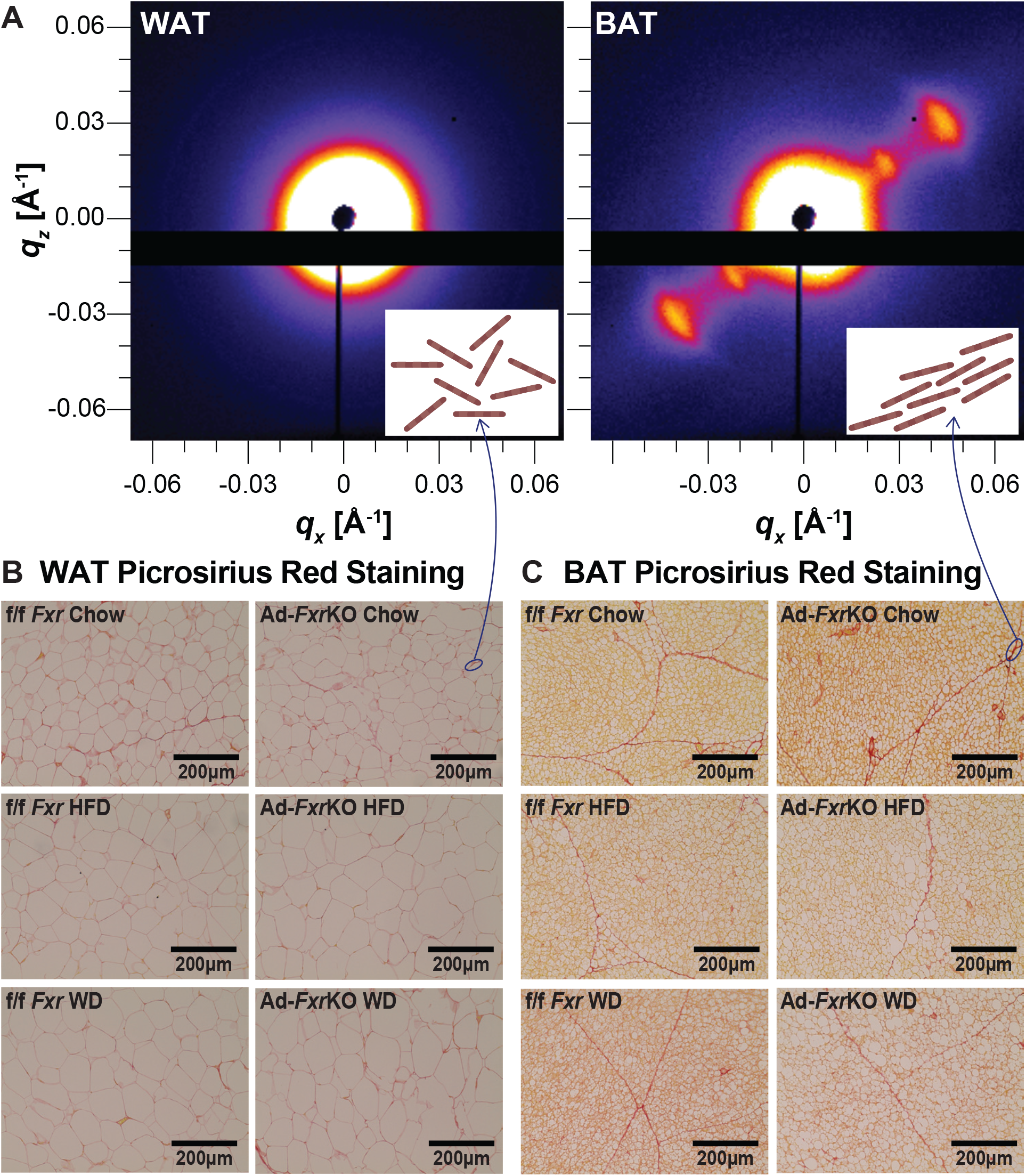
White and brown adipose tissues exhibit distinct collagen orientation. (A) 2D X-ray scattering of BAT and WAT. The inset cartoons represent collagen fibers that are aligned in BAT and randomly oriented in WAT. (B-C) Representative images of picrosirius red-stained WAT (B) and BAT (C) sections of Ad-*Fxr*KO and f/f *Fxr* mice upon chow, HFD, and WD diets for 4 weeks (n=6-8 mice per group).

### HFD and *Fxr* deletion alter BA composition within adipose tissue

BAs, the natural ligands for FXR (*27*), are amphipathic molecules (*30*), and their hydrophilic-hydrophobic ratio determines their capacity to solubilize lipids (*30*). Although hydrophobic BAs increase fat solubility, they are known to be cytotoxic (*31*). We found that f/f *Fxr* BAT showed higher BA levels and lower hydrophobicity compared to WAT (fig. S7, A and B). While Ad-*Fxr*KO mice displayed decreases in β-MCA, the overall hydrophobic index and concentrations remained unaltered compared to f/f *Fxr* mice (Fig. 4, A and B and fig. S7, A and B). We have previously shown the presence of BAs and the expression of genes responsible for BA synthesis and transport in both fat depots (*32*), indicating that depot-specific BA transport and/or local synthesis may occur within adipocytes albeit two orders of magnitude lower than in the liver.

**Figure 4.**
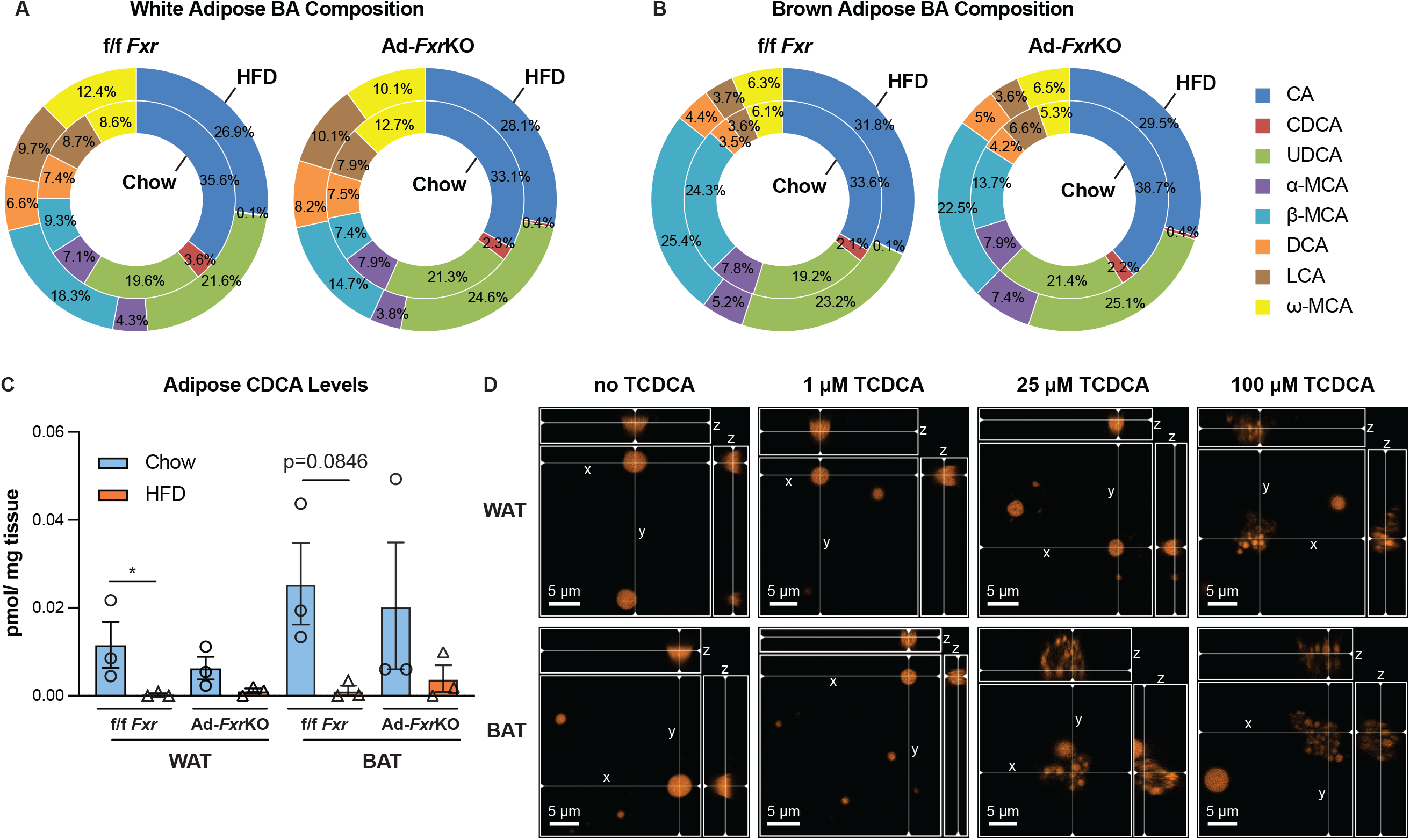
BAs break down lipid droplets, and their composition in the adipose tissue is altered by both HFD and *Fxr* deletion. (A-B) Composition of BAs in the WAT (A) and BAT (B) from Ad-*Fxr*KO and f/f *Fxr* mice upon different diets for 4 weeks (n=3 mice per group). Each BA species includes both free and tauro-conjugated BAs. (C) Levels of TCDCA in the WAT and BAT from Ad-*Fxr*KO and f/f *Fxr* mice upon different diets for 4 weeks (n=3 mice per group). Data are represented as mean ± SEM. \**P* < 0.05. (D) 3D Laser scanning confocal microscopy (LSCM) imaging of isolated LDs from WAT and BAT upon TCDCA treatment. Cross-sections of the LDs are shown on the side for a 3D representation.

HFD-fed f/f *Fxr* mice had different BA composition compared to chow-fed mice in both fat depots (Fig. 4, A and B). For instance, primary BAs that are synthesized in the liver— including cholic acid (CA), chenodeoxycholic acid (CDCA), and α muricholic acid (α-MCA)— were decreased, while primary BA β-MCA and secondary BA ω-MCA that are modified by the gut microbiota were increased in the WAT of HFD-fed compared to the chow-fed f/f *Fxr* mice (Fig. 4A). It is interesting to note that adipose tissue has been shown to harbor microbiome that mimics the gut (*33, 34*) and possibly contributes to the levels of secondary BAs. However, HFD-mediated reduction in CA and increases in β-MCA and ω-MCA were not seen in f/f *Fxr* BAT, suggesting depot-specific alterations to the diet (Fig. 4B). On the other hand, deletion of *Fxr* also leads to alterations in BA composition. Although the ratio of CDCA was lower in the WAT of chow-fed Ad-*Fxr*KO mice, HFD led to a reduction in primary BAs irrespective of the presence or absence of FXR (Fig. 4A). Ad-*Fxr*KO WAT exhibited higher ω-MCA than the f/f *Fxr* mice upon chow, which is reversed under HFD condition (Fig. 4A). In BAT, Ad-*Fxr*KO mice displayed half the amount of β-MCA compared to the f/f *Fxr* mice under chow condition while HFD negated this difference (Fig. 4B). These findings imply that diet and FXR alter primary and secondary BA levels within adipose tissue in a depot specific manner.

### BAs break down LDs

HFD enlarged LD size as anticipated and also led to a robust reduction in primary BAs including CDCA levels in adipose tissue (Fig. 4C). CDCA (*35–37*) has been shown to reduce TAG levels by inhibiting lipogenic genes (*38*), inducing fatty acid β-oxidation genes (*39*), and clearing TAGs in an FXR-dependent manner (*40*) or promote energy expenditure via the membrane receptor TGR5 (*41*). We examined if BAs, besides initiating an intracellular signaling cascade, have a direct role as natural surfactants in regulating LD physical properties in adipose tissue. We isolated LDs and imaged them at different stacked focal planes in the presence and absence of taurochenodeoxycholic acid (TCDCA), the major form of CDCA in mice, using laser scanning confocal microscopy (LSCM). TCDCA at 25 μM or 100 μM—concentrations that are much lower than their critical micellar concentration (CMC) of 3 mM (*42*)—were sufficient to disintegrate LDs obtained from BAT and WAT into smaller LDs (Fig. 4D). This result suggests that TCDCA can stimulate the breakdown of large LDs and may stabilize small LDs. Combining our data together, we postulate that HFD-induced reduction in TCDCA in adipose tissue may promote LDs with more fat layers during caloric excess (Fig. 2C). Unmicellized BA surfactants are soluble in the cytosol but tend to adsorb at the LD oil-water interface, reducing its surface tension and facilitating the formation of smaller LDs, which is a putative mechanism underlying LD fission during lipolysis (*43*).

Overall, our findings reveal that deletion of adipose FXR alters local BA composition and adipose expansion. We show that, beyond size, LDs regulate their structure and fat packing during obesity and that adipose depot-specific collagen orientation may regulate and constrain the extent of adipose tissue expansion. Finally, we demonstrate that BAs in adipose tissue can remodel LD size.

## Acknowledgments

This work was supported by the Cancer Center at Illinois (FY21, CL and SB), the National Institutes of Health Grant No. 1DP2EB024377-01 (CL and SB), the National Institute of Diabetes and Digestive and Kidney Diseases, R01 DK113080 (SA), USDA HATCH funds ILLU-971-377 (SA), American Cancer Society Grant RSG-132315 (SA), Cancer center at Illinois (SA) and the Office of Naval Research (ONR) N000141812087 DURIP—Defense University Research Instrumentation Program (LCSM, CL and SB). SAXS experiments were carried out at the beamline 12-ID-B at the Advanced Photon Source (APS), Argonne National Laboratory. Use of APS was supported by the US Department of Energy (DOE), Office of Science, Office of Basic Energy Sciences, under contract number DE-AC02-06CH11357. Adipose tissue BA analysis was performed at the NIH West Coast Metabolomics Center at the University of California, Davis.

## Author contributions

W.Z., S.R.B., C.L. and S.A. conceived and designed research; W.Z. and S.R.B. performed experiments and analyzed data; W.Z., S.R.B., C.L. and S.A interpreted data; W.Z. and S.R.B generated figures and drafted the manuscript; all authors were involved in editing and revising the manuscript, and had final approval of the submitted and published versions.

## Data and materials availability

All data are in the main text and the supplementary materials.

