## Supplementary Materials for "Lipid droplet structural remodeling in adipose tissue upon caloric excess"

**The PDF file includes:**

Materials and Methods

Figs. S1 to S7

Tables S1 to S3

References

### Materials and Methods

#### Animals

Floxed *Fxr* (f/f *Fxr*) mice obtained from Dr. Kristina Schoonjans at Ecole Polytechnique Federale de Lausanne, were bred with mice expressing the Cre-recombinase under the control of adiponectin promoter (Adipoq-Cre, Jackson Laboratory) to generate adipocyte-specific *Fxr* knockout (Ad- *Fxr*KO) mice. Male Ad- *Fxr*KO and f/f *Fxr* mice at 10- to 12-week-old were used. These mice were bred and maintained on a 12:12 light/dark cycle with *ad libitum* access to tap water and a normal chow diet in a climate-controlled (23 °C) animal facility at the University of Illinois at Urbana-Champaign. At 10-12 weeks, mice were fed a normal chow (Envigo), 60% high fat (Envigo), or high fat/high sucrose western (Envigo) diet for 4 weeks to mimic obesogenic conditions. Mice were weighed weekly. After 4 weeks, mice were sacrificed at the end of the experimental regimen. Interscapular brown (BAT) and gonadal white (WAT) adipose tissues were collected for isolation of mature adipocytes and analysis of histology, gene expression, fat packing, and bile acid (BA) levels. To examine *Fxr* transcript levels in isolated mature adipocytes and the role of BAs in lipid droplet (LD) structural properties, male C57BL/6 wild-type (WT) mice at 3- to 4-week-old were sacrificed. BAT and WAT were collected for mature adipocytes and LD isolation. All experiments were performed following the National Institutes of Health guidelines for the care and use of laboratory animals, and all procedures were approved by the Institutional Animal Care and Use Committee at the University of Illinois at Urbana-Champaign.

#### Isolation of mature adipocytes

Mature adipocytes were isolated from Interscapular brown and inguinal white adipose tissue as described previously (*1-3*). Briefly, male C57BL/6 WT mice at 3- to 4-week-old were euthanized by isoflurane inhalational anesthesia followed by cervical dislocation. The interscapular brown and inguinal white adipose depots were harvested and minced with scissors in DMEM (Gibco). Tissue fragments were incubated with digestion buffer ((dd$H_2O$ containing HEPES (100 mM;

Fisher Bioreagents), NaCl (123 mM; Fisher Chemical), KCl (5 mM; Fisher Chemical), $CaCl_2$ (1.3 mM; Fisher Chemical), glucose (5 mM; Fisher Chemical), bovine serum albumin (BSA) (1.5% w/v; VWR) and collagenase type I (2 mg/mL; Worthington Biochemical Corporation)), and shaken at 300 rpm at 37 °C for 1 hour. The digested solution was then passed through a 100-µm cell strainer and placed on ice for 20 min. The infranatant below the top mature adipocyte layer was removed, and the mature adipocytes were washed with DMEM (Gibco) followed by centrifugation at 200 g for 5 min. Then the top mature adipocyte layer was collected and lysed in TRIzol reagent (Invitrogen) for the following gene expression analysis.

**Quantitative real-time PCR**

Total RNA was isolated from mature adipocytes and snap-frozen adipose tissues using TRIzol reagent (Invitrogen). Upon DNase I (New England Biolabs) treatment, RNA was reverse transcribed into cDNA using a Maxima reverse transcriptase kit (Thermo Scientific). Quantitative real-time PCR (qRT-PCR) was performed with SYBR green master mix (Applied Biosystems) using Applied Biosystems QuantStudio 7 Flex Real-Time PCR System. To determine relative expression values, the $2^{-\Delta\Delta Ct}$ method was used, where triplicate Ct values for each sample were averaged and subtracted from those derived from housekeeping gene *36B4*. All primers used are listed in Table S1.

**Histology**

Adipose tissues were fixed in 10% neutral-buffered formalin (VWR) for 24 hours at 4 °C and processed. Formalin-fixed tissues were embedded using paraffin and cut on a microtome at 5 µm thickness and mounted onto charged glass microscope slides. Adipose sections were deparaffinized and stained with hematoxylin & eosin (Thermo Scientific) using standard histological protocol. Adipocyte size was quantified using Adiposoft-ImageJ software. Collagen was stained using picrosirius red (Sigma-Aldrich).

**Small-angle X-ray scattering (SAXS) of adipose tissue**

SAXS experiments were carried out at Beamline 12-ID-B at the Advanced Photon Source at Argonne National Laboratory. An average photon energy of 13.3 keV was used. Tissue samples were either loaded onto a solid sample holder using Kapton tape or centrifuged into quartz capillaries. 2D scattering data were radially averaged using IGOR Pro. The q-calibrant was silver behenate. Data analysis was carried out using Mathematica.

**Bile acid analysis**

Adipose bile acid (BA) analysis was performed in the NIH West Coast Metabolomics Center at the University of California, Davis. Adipose BAs were extracted from interscapular brown and gonadal white adipose tissue samples (4-4.75 mg) as previously described (*4*). Six internal standards (GCA-d4, TCDCA-d4, CA-d6, GCDCA-d4, CDCA-d4, DCA-d4) were added. BA levels were quantified by ultra-high performance liquid chromatography chromatography-triple quadruple mass spectrometry (UHPLC-TQ-MS/MS) (Thermo Fisher Scientific).

**Laser scanning confocal microscopy of isolated LDs**

LDs were isolated from brown adipose tissue (BAT) and white adipose tissue (WAT) of chow-fed mice based on density gradient centrifugation as described previously (*5*). The isolated LDs were incubated with Droplite™ Red staining solution (AAT Bioquest, Cat # 22735). 50 µL of Droplite™ Red staining solution was added to 100 µL of each of the LD samples, and the samples were incubated at 37 °C for 30 minutes. Bile acid TCDCA was then added to the stained LD solution to achieve a final concentration of 1 µM, 25 µM, or 100 µM, and incubated at room temperature for at least 1.5 hours. The samples were then imaged using an LSM800 confocal microscope with a 63x lens at the Leal Lab at UIUC. An excitation wavelength of 561 nm was used for the Droplite™ dye.

**Statistical analysis**

Data were expressed as means ± SEM. Statistical analyses were performed using GraphPad Prism 8 software. Differences between two groups were analyzed using Student's *t* test, and multiple group comparisons were analyzed using a two-way or three-way ANOVA with a Fisher's LSD *post hoc* test. $P < 0.05$ was considered statistically significant.

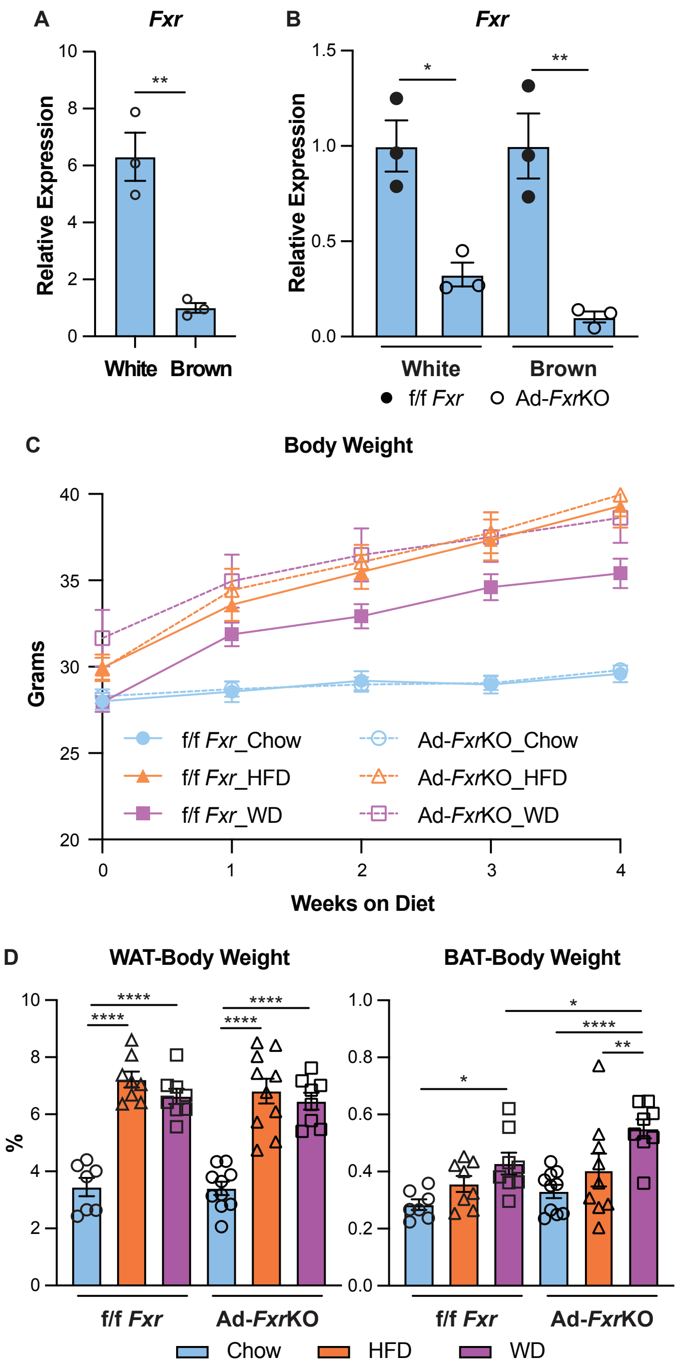


**Supplemental Figure 1. Adipocyte-specific knockout of *Fxr* does not affect body weight or white fat mass under either normal or obese condition.** (A) *Fxr* mRNA levels in mature adipocytes isolated from brown (BAT) and gonadal white (WAT) adipose tissue from wild-type mice (n=3 mice per group). (B) *Fxr* mRNA levels in mature adipocytes isolated from BAT and WAT of adipocyte-specific *Fxr* knockout (Ad-*Fxr*KO) and f/f *Fxr* control mice (n=3 mice per group). (C) Weekly body weight of Ad-*Fxr*KO and f/f *Fxr* mice fed with either normal chow, 60% high fat (HFD), or high fat/high sucrose western (WD) diet for 4 weeks (n=8-10 mice per group). (D) WAT or BAT to body weight ratios after 4 weeks of diet (n=7-10 mice per group). Data are represented as mean ± SEM. *$P < 0.05$, **$P < 0.01$, ****$P < 0.0001$.

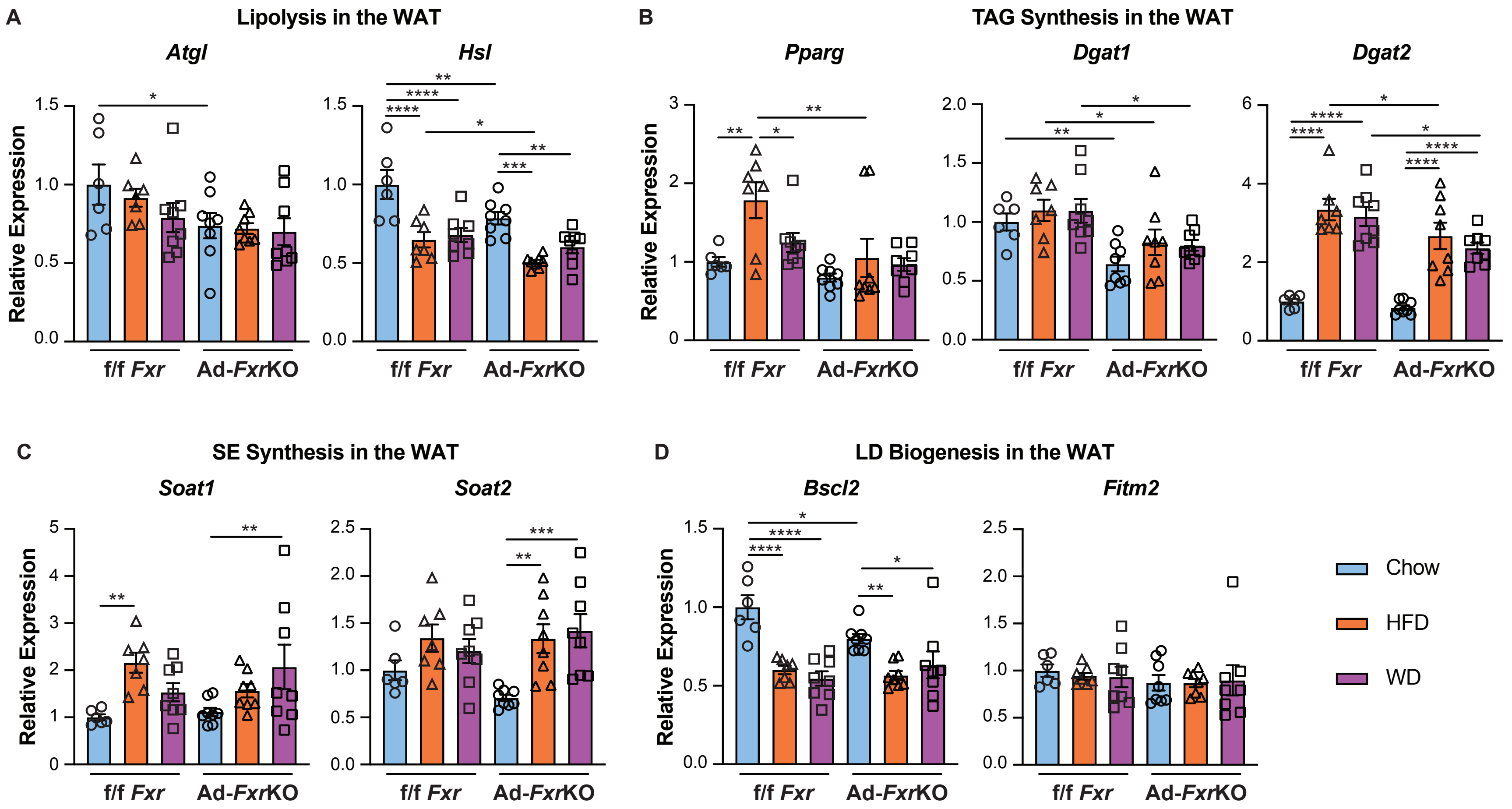


**Supplemental Figure 2. Obesogenic diets and *Fxr* deletion lead to altered expression of genes involved in LD breakdown and synthesis in the WAT.** (A-D) mRNA levels of genes related to lipolysis (A), TAG (B) and SE (C) synthesis, and LD biogenesis (D) in the WAT from Ad-*Fxr*KO and f/f *Fxr* mice upon different diets for 4 weeks (n=6-8 mice per group). Data are represented as mean ± SEM. **P* < 0.05, ***P* < 0.01, ****P* < 0.001, *****P* < 0.0001.

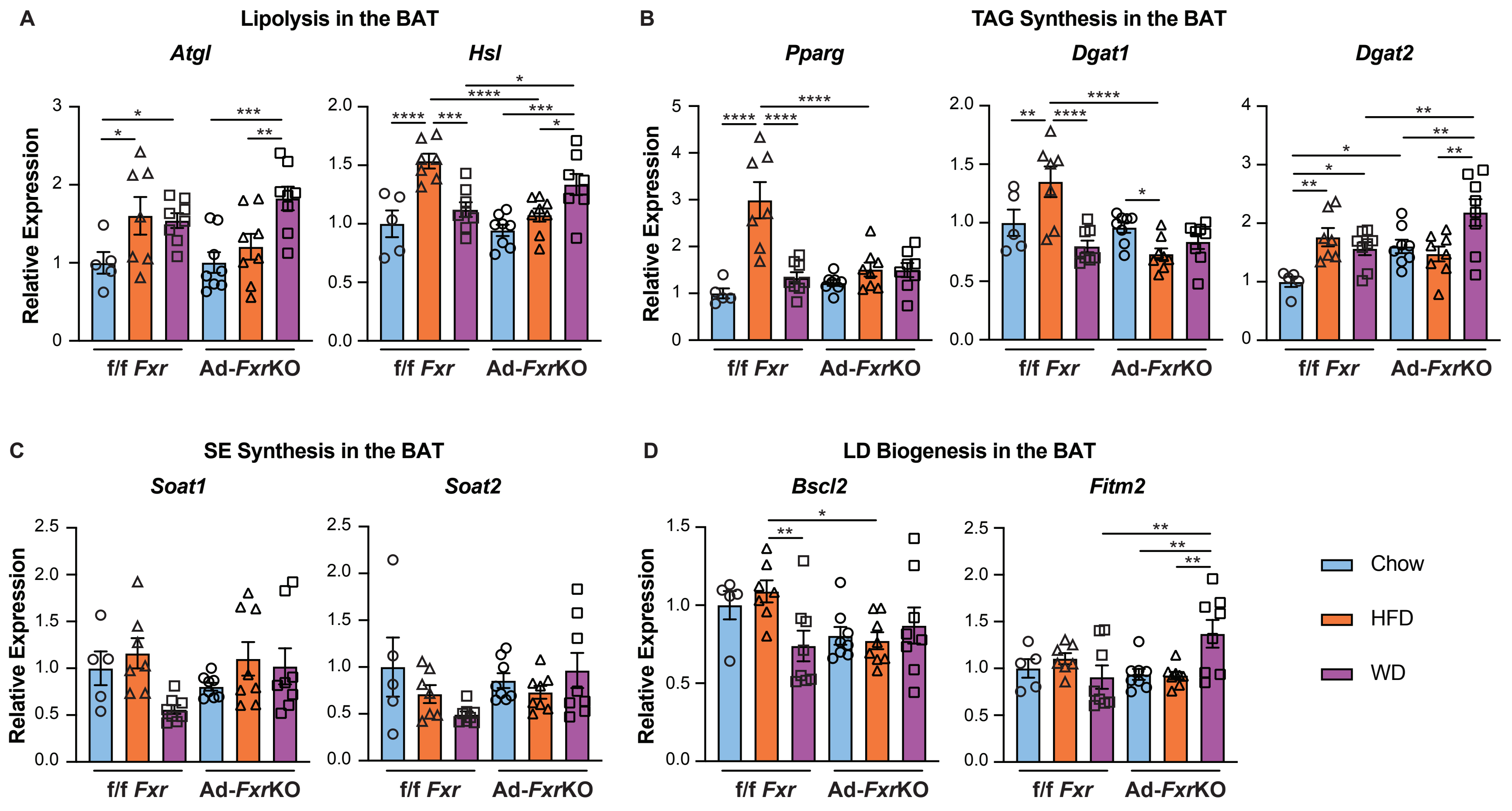


**Supplemental Figure 3. Obesogenic diets and *Fxr* knockout result in changes in the expression of genes related to LD biosynthesis and breakdown in the BAT.** (A-D) mRNA levels of genes related to lipolysis (A), TAG (B) and SE (C) synthesis, and LD biogenesis (D) in the WAT from Ad-*Fxr*KO and f/f *Fxr* mice upon different diets for 4 weeks (n=5-8 mice per group). Data are represented as mean ± SEM. **P* < 0.05, ***P* < 0.01, ****P* < 0.001, *****P* < 0.0001.

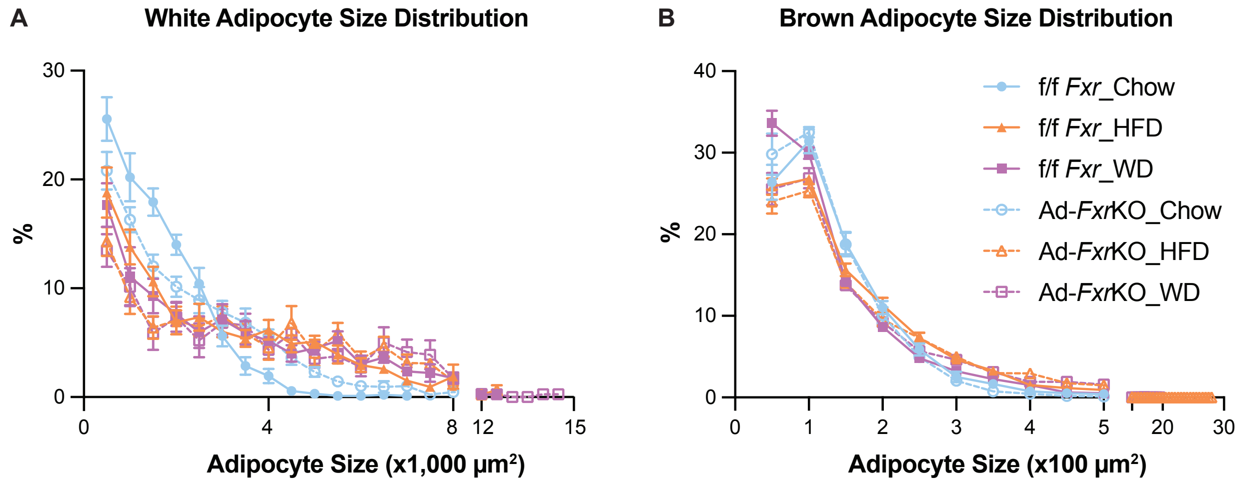


**Supplemental Figure 4. *Fxr* knockout causes adipocyte hypertrophy.** (A-B) Distribution of adipocyte size of H&E-stained WAT (A) and BAT (B) sections of Ad-*Fxr*KO and f/f *Fxr* mice upon different diets for 4 weeks (n=6-8 mice per group).

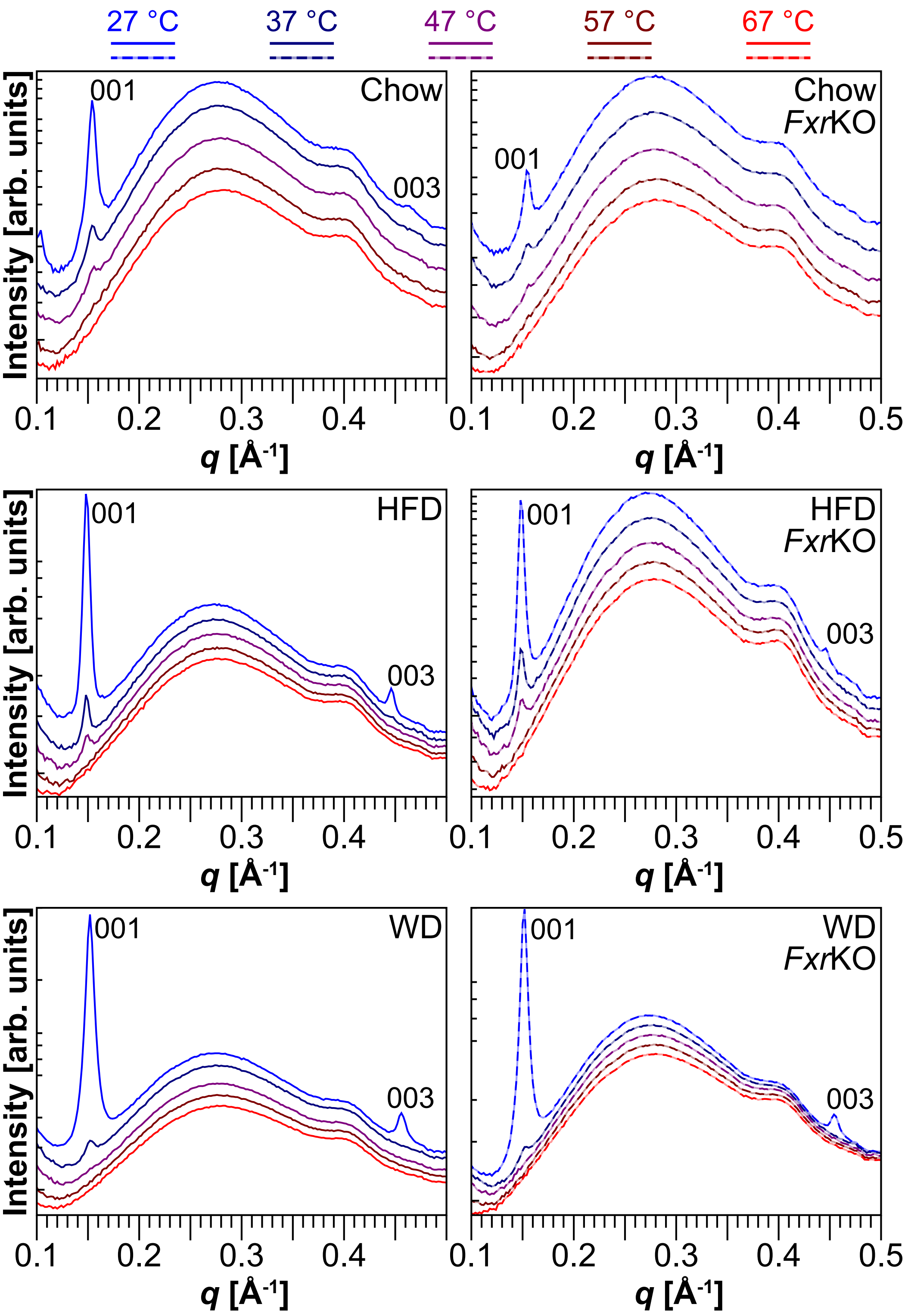


**Supplemental Figure 5. Increasing temperature decreases multilamellar peak intensity.** Increasing the temperature from 27 °C to 67 °C reduces the intensities of the 001 and 003 multilamellar peaks in WAT (n=1 mouse per group, 7 locations on tissue per mouse). The broad peak at around 0.4 Å$^{-1}$ is an artifact of the holder.

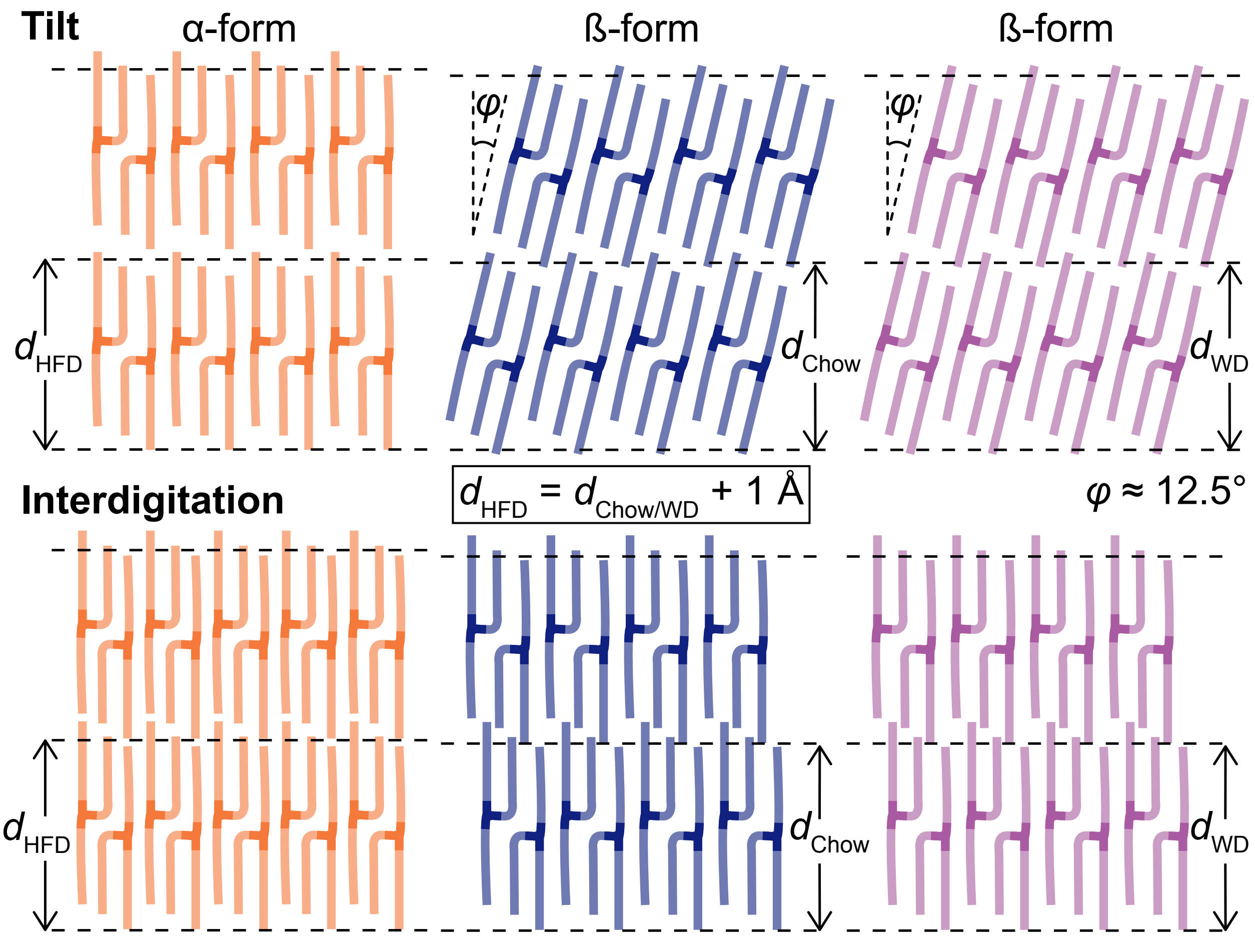


**Supplemental Figure 6. TAGs at the multilamellar layers could assume either tilted or interdigitated conformations.** Tilt or interdigitation could account for the consistent 1 Å difference in repeat spacing between HFD and chow/WD diets (however, no difference is observed in WAXS peak positions between different diets).

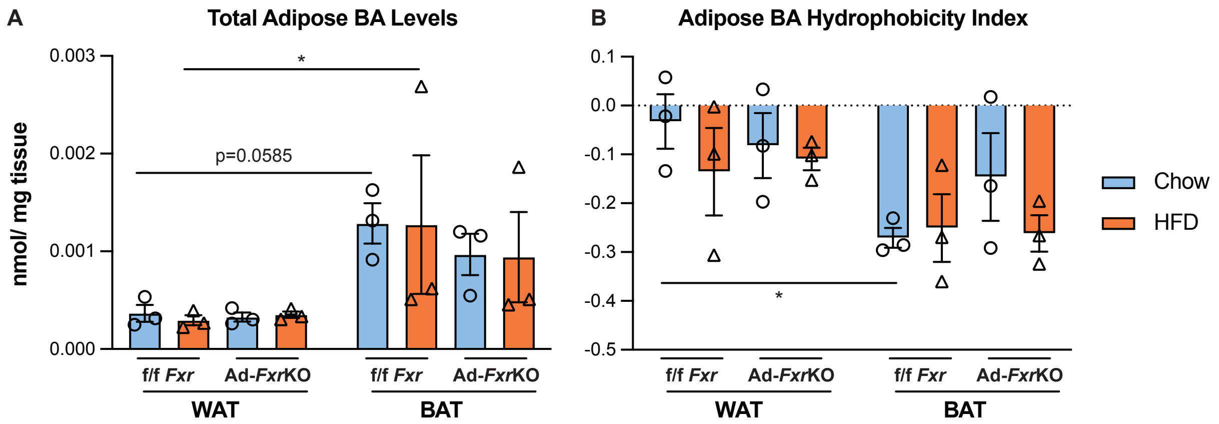


| | |
|---|---|
| Depot | p=0.0032 |
| Diet | p=0.9244 |
| Genotype | p=0.4954 |
| Depot x Diet | p=0.9935 |
| Depot x Genotype | p=0.4700 |
| Diet x Genotype | p=0.9297 |
| Depot x Diet x Genotype | p=0.9046 |

| | |
|---|---|
| Depot | p=0.0050 |
| Diet | p=0.2157 |
| Genotype | p=0.6138 |
| Depot x Diet | p=0.8445 |
| Depot x Genotype | p=0.4469 |
| Diet x Genotype | p=0.7351 |
| Depot x Diet x Genotype | p=0.2457 |

**Supplemental Figure 7. BAT exhibits higher BA levels but lower hydrophobicity than WAT.** (A-B) Levels (A) and hydrophobicity index (B) of BAs in the WAT and BAT from Ad-*Fxr*KO and f/f *Fxr* mice upon different diets for 4 weeks (n=3 mice per group).

**Supplementary Table 1. Primer sequences used in qRT-PCR analysis**

| Gene | Forward (5' -> 3') | Reverse (5' -> 3') |
|---|---|---|
| *36b4* | AGATGCAGCAGATCCGCAT | GTTCTTGCCCATCAGCACC |
| *Fxr* | ACAGCTAATGAGGACGACAG | GATTTCCTGAGGCATTCTCTG |
| *Atgl* | TCCGTGGCTGTCTACTAAAGA | TGGGATATGATGACGTTCTCTCC |
| *Hsl* | CTGGTGCAGAGAGACACTTC | CTTGCGTCCACTTAGTTCCA |
| *Pparg* | CAAGAATACCAAAGTGCGATCAA | GAGCTGGGTCTTTTCAGAATAATAAG |
| *Dgat1* | CAGCTGTGGCCTTACTGGTT | AGAGACAGCTTTGGCCTTGA |
| *Dgat2* | CTCTGTCACCTGGCTCAACA | TATCAGCCAGCAGTCTGTGC |
| *Soat1* | GAAACCGGCTGTCAAAATCTGG | TGTGACCATTTCTGTATGTGTCC |
| *Soat2* | TCACCATGAACGACAGGCAC | GAATGTTGTCTGGGGCAGGG |
| *Bscl2* | TGGGGCAAGAGAGACATGC | TCTTCCACAGGGACGATACCC |
| *Fitm2* | TCGGTCGTCAAGGAGCTGT | CAAAATACACGTTGAGGACGTTG |

**Supplementary Table 2. WAT bile acid concentrations in Ad-*Fxr*KO and f/f *Fxr* mice upon chow and HFD**

| Unit: nmol/mg tissue | CA | TCA | TCDCA | UDCA | TUDCA | α-MCA | T-α-MCA | β-MCA |
|---|---|---|---|---|---|---|---|---|
| **Ad-*Fxr*KO-Chow-WAT-1** | 1.20E-04 | 2.83E-05 | 1.12E-05 | 6.25E-05 | 9.47E-06 | 2.65E-05 | 6.67E-06 | 2.70E-06 |
| **Ad-*Fxr*KO-Chow-WAT-2** | 3.32E-05 | 3.57E-05 | 2.35E-06 | 2.93E-05 | 1.15E-05 | 9.08E-06 | 1.80E-05 | 3.07E-06 |
| **Ad-*Fxr*KO-Chow-WAT-3** | 3.43E-05 | 2.20E-05 | 5.44E-06 | 5.77E-05 | 5.42E-06 | 3.41E-06 | 1.96E-06 | 5.08E-06 |
| **f/f *Fxr*-Chow-WAT-1** | 5.36E-05 | 3.04E-05 | 8.53E-06 | 4.52E-05 | 6.46E-06 | 1.23E-05 | 1.09E-05 | 6.38E-06 |
| **f/f *Fxr*-Chow-WAT-2** | 1.24E-04 | 5.96E-05 | 2.17E-05 | 6.12E-05 | 3.51E-05 | 2.00E-05 | 1.72E-05 | 7.83E-06 |
| **f/f *Fxr*-Chow-WAT-3** | 4.96E-05 | 2.35E-05 | 4.48E-06 | 3.76E-05 | 2.41E-06 | 4.89E-06 | 2.64E-06 | 0.00E+00 |
| **Ad-*Fxr*KO-HFD-WAT-1** | 2.30E-05 | 4.68E-05 | 0.00E+00 | 2.66E-05 | 5.70E-05 | 0.00E+00 | 7.91E-06 | 1.19E-05 |
| **Ad-*Fxr*KO-HFD-WAT-2** | 3.40E-05 | 6.42E-05 | 2.03E-06 | 3.32E-05 | 4.38E-05 | 3.88E-06 | 1.48E-05 | 6.60E-06 |
| **Ad-*Fxr*KO-HFD-WAT-3** | 2.44E-05 | 5.95E-05 | 1.16E-06 | 2.82E-05 | 3.14E-05 | 1.08E-06 | 6.77E-06 | 3.49E-06 |
| **f/f *Fxr*-HFD-WAT-1** | 2.12E-05 | 6.69E-05 | 0.00E+00 | 1.98E-05 | 4.59E-05 | 0.00E+00 | 2.07E-05 | 3.88E-06 |
| **f/f *Fxr*-HFD-WAT-2** | 1.70E-05 | 4.60E-05 | 0.00E+00 | 1.86E-05 | 2.98E-05 | 0.00E+00 | 7.10E-06 | 2.70E-06 |
| **f/f *Fxr*-HFD-WAT-3** | 2.25E-05 | 2.59E-05 | 8.83E-07 | 2.92E-05 | 1.69E-05 | 0.00E+00 | 4.03E-06 | 3.56E-06 |

| | T-β-MCA | DCA | TDCA | LCA | TLCA | ω-MCA | T-ω-MCA |
|---|---|---|---|---|---|---|---|
| **Ad-*Fxr*KO-Chow-WAT-1** | 1.9E-05 | 1.74E-05 | 0 | 3.41E-06 | 2.25E-05 | 1.85E-05 | 1.99E-05 |
| **Ad-*Fxr*KO-Chow-WAT-2** | 2.84E-05 | 1.58E-05 | 4.13E-06 | 2.64E-06 | 1.37E-05 | 1.78E-05 | 2.85E-05 |
| **Ad-*Fxr*KO-Chow-WAT-3** | 2.66E-06 | 1.77E-05 | 6.52E-06 | 7.19E-06 | 1.56E-05 | 1.76E-05 | 2.51E-06 |
| **f/f *Fxr*-Chow-WAT-1** | 1.77E-05 | 1.62E-05 | 5.22E-06 | 1.36E-05 | 1.72E-05 | 1.76E-05 | 3.50E-06 |
| **f/f *Fxr*-Chow-WAT-2** | 5.24E-05 | 1.44E-05 | 8.17E-06 | 3.25E-06 | 2.51E-05 | 1.65E-05 | 2.33E-05 |
| **f/f *Fxr*-Chow-WAT-3** | 4.49E-06 | 2.22E-05 | 4.44E-06 | 2.24E-06 | 2.21E-05 | 1.73E-05 | 4.45E-06 |
| **Ad-*Fxr*KO-HFD-WAT-1** | 2.28E-05 | 1.09E-05 | 7.35E-06 | 9.75E-06 | 1.26E-05 | 1.60E-05 | 8.63E-06 |
| **Ad-*Fxr*KO-HFD-WAT-2** | 5.17E-05 | 2.20E-05 | 8.89E-06 | 2.35E-05 | 1.70E-05 | 1.78E-05 | 1.74E-05 |
| **Ad-*Fxr*KO-HFD-WAT-3** | 3.58E-05 | 1.23E-05 | 1.22E-05 | 1.40E-05 | 1.41E-05 | 1.91E-05 | 1.12E-05 |
| **f/f *Fxr*-HFD-WAT-1** | 8.74E-05 | 1.13E-05 | 6.40E-06 | 5.44E-06 | 1.38E-05 | 1.64E-05 | 3.14E-05 |
| **f/f *Fxr*-HFD-WAT-2** | 2.94E-05 | 1.07E-05 | 8.33E-06 | 2.16E-06 | 2.37E-05 | 1.63E-05 | 8.01E-06 |
| **f/f *Fxr*-HFD-WAT-3** | 8.98E-06 | 1.24E-05 | 0.00E+00 | 1.06E-05 | 1.64E-05 | 1.82E-05 | 1.89E-06 |

**Supplementary Table 3. BAT bile acid concentrations in Ad-*Fxr*KO and f/f *Fxr* mice upon chow and HFD**

| Unit: nmol/mg tissue | CA | TCA | TCDCA | UDCA | TUDCA | α-MCA | T-α-MCA | β-MCA |
|---|---|---|---|---|---|---|---|---|
| **Ad-*Fxr*KO-Chow-BAT-1** | 4.93E-04 | 3.67E-05 | 4.93E-05 | 2.17E-04 | 2.24E-05 | 1.09E-04 | 9.86E-06 | 1.37E-05 |
| **Ad-*Fxr*KO-Chow-BAT-2** | 8.83E-05 | 7.44E-05 | 6.02E-06 | 7.65E-05 | 4.60E-05 | 1.96E-05 | 1.57E-05 | 2.68E-05 |
| **Ad-*Fxr*KO-Chow-BAT-3** | 6.11E-05 | 3.14E-04 | 6.04E-06 | 6.66E-05 | 1.60E-04 | 7.93E-06 | 5.43E-05 | 5.30E-05 |
| **f/f *Fxr*-Chow-BAT-1** | 1.50E-04 | 2.58E-04 | 1.94E-05 | 8.51E-05 | 1.45E-04 | 2.20E-05 | 5.60E-05 | 3.82E-05 |
| **f/f *Fxr*-Chow-BAT-2** | 2.61E-04 | 2.75E-04 | 4.37E-05 | 1.30E-04 | 1.45E-04 | 4.60E-05 | 1.14E-04 | 4.59E-05 |
| **f/f *Fxr*-Chow-BAT-3** | 1.09E-04 | 1.90E-04 | 1.34E-05 | 7.61E-05 | 1.30E-04 | 1.00E-05 | 4.07E-05 | 2.25E-05 |
| **Ad-FxrKO-HFD-BAT-1** | 4.53E-05 | 7.43E-05 | 1.83E-06 | 4.04E-05 | 1.01E-04 | 1.03E-06 | 2.04E-05 | 1.67E-05 |
| **Ad-FxrKO-HFD-BAT-2** | 1.38E-04 | 4.17E-04 | 9.93E-06 | 9.34E-05 | 3.14E-04 | 1.88E-05 | 1.31E-04 | 7.14E-05 |
| **Ad-FxrKO-HFD-BAT-3** | 2.81E-05 | 8.70E-05 | 0.00E+00 | 3.84E-05 | 8.67E-05 | 0.00E+00 | 2.62E-05 | 1.18E-05 |
| **f/f *Fxr*-HFD-BAT-1** | 3.44E-05 | 8.69E-05 | 2.01E-07 | 3.66E-05 | 8.76E-05 | 0.00E+00 | 1.85E-05 | 1.71E-05 |
| **f/f *Fxr*-HFD-BAT-2** | 3.21E-05 | 8.39E-04 | 0.00E+00 | 2.79E-05 | 5.57E-04 | 1.69E-06 | 1.50E-04 | 7.31E-05 |
| **f/f *Fxr*-HFD-BAT-3** | 6.40E-05 | 1.09E-04 | 3.45E-06 | 4.31E-05 | 9.78E-05 | 1.77E-06 | 2.07E-05 | 1.95E-05 |

| | T-β-MCA | DCA | TDCA | LCA | TLCA | ω-MCA | T-ω-MCA |
|---|---|---|---|---|---|---|---|
| **Ad-*Fxr*KO-Chow-BAT-1** | 2.29E-05 | 2.63E-05 | 5.70E-06 | 1.04E-05 | 1.10E-04 | 1.69E-05 | 9.94E-06 |
| **Ad-*Fxr*KO-Chow-BAT-2** | 5.02E-05 | 1.47E-05 | 1.30E-05 | 1.52E-05 | 1.42E-05 | 1.88E-05 | 1.49E-05 |
| **Ad-*Fxr*KO-Chow-BAT-3** | 2.12E-04 | 1.18E-05 | 4.36E-05 | 1.24E-05 | 1.85E-05 | 1.74E-05 | 6.89E-05 |
| **f/f *Fxr*-Chow-BAT-1** | 3.33E-04 | 1.82E-05 | 1.86E-05 | 3.75E-05 | 1.65E-05 | 1.69E-05 | 5.36E-05 |
| **f/f *Fxr*-Chow-BAT-2** | 3.34E-04 | 2.11E-05 | 2.68E-05 | 1.06E-05 | 3.41E-05 | 1.88E-05 | 6.84E-05 |
| **f/f *Fxr*-Chow-BAT-3** | 1.26E-04 | 1.94E-05 | 2.40E-05 | 1.75E-05 | 1.65E-05 | 1.85E-05 | 4.92E-05 |
| **Ad-FxrKO-HFD-BAT-1** | 4.02E-05 | 1.21E-05 | 0.00E+00 | 1.72E-05 | 1.52E-05 | 1.65E-05 | 7.80E-06 |
| **Ad-FxrKO-HFD-BAT-2** | 3.73E-04 | 2.90E-05 | 5.94E-05 | 2.87E-05 | 1.55E-05 | 1.64E-05 | 9.63E-05 |
| **Ad-FxrKO-HFD-BAT-3** | 9.02E-05 | 1.88E-05 | 1.41E-05 | 3.46E-06 | 1.66E-05 | 1.77E-05 | 1.97E-05 |
| **f/f *Fxr*-HFD-BAT-1** | 9.96E-05 | 1.51E-05 | 1.81E-06 | 1.10E-05 | 1.46E-05 | 1.89E-05 | 1.56E-05 |
| **f/f *Fxr*-HFD-BAT-2** | 6.43E-04 | 1.81E-05 | 8.18E-05 | 3.41E-05 | 1.52E-05 | 1.86E-05 | 1.45E-04 |
| **f/f *Fxr*-HFD-BAT-3** | 7.93E-05 | 2.14E-05 | 2.33E-05 | 1.81E-05 | 4.09E-05 | 1.65E-05 | 1.56E-05 |
